# Autologous adipose-derived stem cell transplantation enhances healing of wound with exposed bone in a rat model

**DOI:** 10.1101/571968

**Authors:** Tomo Hamada, Hidenori Matsubara, Yasuhisa Yoshida, Shuhei Ugaji, Issei Nomura, Hiroyuki T suchiya

## Abstract

**Objectives:** Soft tissue wounds with exposed bone often require extended healing times and can be associated with severe complications. We describe the ability of artificial dermis with autogenic adipose-derived stem cells (ADSCs) to promote the healing of wounds with exposed bone in a rat model.

**Methods:** Adipose tissues harvested from the bilateral inguinal regions of Wistar rats were used as ADSCs. Rats were randomly divided into control and ADSC groups to investigate the efficacy of ADSC transplantation for wound healing (n=20 per group). Soft tissue defects were created on the heads of the rats and were covered with artificial dermis with or without the seeded ADSCs. Specimens from these rats were evaluated using digital image analysis, histology, immunohistochemistry, cell labeling, and real-time reverse-transcription polymerase chain reaction (Real-time RT-PCR).

**Results:** The average global wound area was significantly smaller in the ADSC group than in the control group on days 3, 7, and 14 after surgery (*p*<0.05). After 14 days, the blood vessel density in the wound increased by 1.6-fold in the ADSC group compared with that in the control group (*p*<0.01). Real-time RT-PCR results showed higher *Fgfb* and *Vegf* expression levels at all time points, and higher *Tgfb1* and *Tgfb3* expression levels until 14 days after surgery, in the ADSC group than in the control group (*p*<0.05).

**Conclusions:** In wounds with exposed bone, autogenic ADSCs can promote vascularization and wound healing. Use of this cell source has multiple benefits, including convenient clinical application and lack of ethical concerns.

## Introduction

Some wounds caused by ulcers, trauma, and various operations, result in exposed bone, leading to severe complications in many cases during treatment of soft tissue. Currently, surgical treatment for defects with exposed bone typically involves the use of local or distal skin flaps, muscle flaps, or myocutaneous flaps. However, there are many risks associated with wound coverage by these flaps, and a complicated microsurgical approach is required for successful treatment. Moreover, use of a composite tissue transfer technique may not be possible in such cases owing to the paucity of the graft donor site and other factors.

Artificial dermis is a commercially available treatment for full-thickness skin defects after debridement. It has been successfully used to promote healing by creating a vascular matrix over an exposed bone in clinical reports [1,2]. However, the formation of neodermal tissue is delayed, thus prolonging the treatment period. A main reason for prolonging the treatment period is the slow vascularization rate [3]. Several recent studies have shown that in the context of tissue injury, mesenchymal stem cells (MSCs) exhibit excellent potential for promoting healing and vascularization of wounds [4,5]. Among the various types of MSCs, adipose-derived stromal stem cells (ADSCs) have many unique advantages. ADSCs are abundant in the subcutaneous adipose tissue and can be easily harvested using a syringe or a minimally invasive lipoaspiration procedure.

Our aim was to evaluate the efficacy of autogenic ADSCs with artificial dermis to promote the healing and vascularization of a wound with exposed bone, which is one of the wounds with the worst healing conditions.

## Methods

### Animal experiments

All experimental protocols were approved by the Kanazawa University Advanced Science Research Center (Approval Number: AP-173885). Forty-three female Wistar rats [age, 8-9 weeks and mean weight, 147.4 g (± 9.3 g)] were used in this experiment, and housed under specific pathogen-free conditions with three rats per cage in a 12-hour light/dark cycle with ad libitum access to food and water. A schematic overview of the experimental design is shown in Fig. 1. The rats were anesthetized with an intraperitoneal injection of pentobarbital (40 mg/kg) and xylazin (15 mg/kg). The adipose tissues were harvested from the bilateral inguinal region (total yield of approximately 1.2 g) for use as ADSCs. After one week, 12 × 12-mm^2^ circular soft tissue defects with exposed bone were created on the heads of the rats by removing the cutaneous tissue and the periosteum of the cranium using a modification of a previously reported method [6]. After creating the soft tissue defect, all rats were housed with one rat per cage. The rats were randomly divided into two groups (a control group and an ADSC group, 20 rats each) to investigate the efficacy of ADSC transplantation for wound healing. The soft tissue defects were then covered with a 12 × 12-mm^2^ circular artificial dermis with or without the seeded ADSCs. Defects were closed with six stitches using 5-0 monofilament nylon sutures (Keisei Medical, Tokyo, Japan), and the extent of wound healing of the defects was observed at 3, 7, 14, and 21 days after surgery (n = 5 at each time point). All rats from each group were euthanized after tracing the wound area on the photograph taken at the established endpoint. The specimens from these rats were then evaluated using digital image analysis, histology, immunohistochemistry, and real-time reverse-transcription-polymerase chain reaction (RT-PCR). The three remaining rats were used for DiI labeling.

**Fig. 1.**
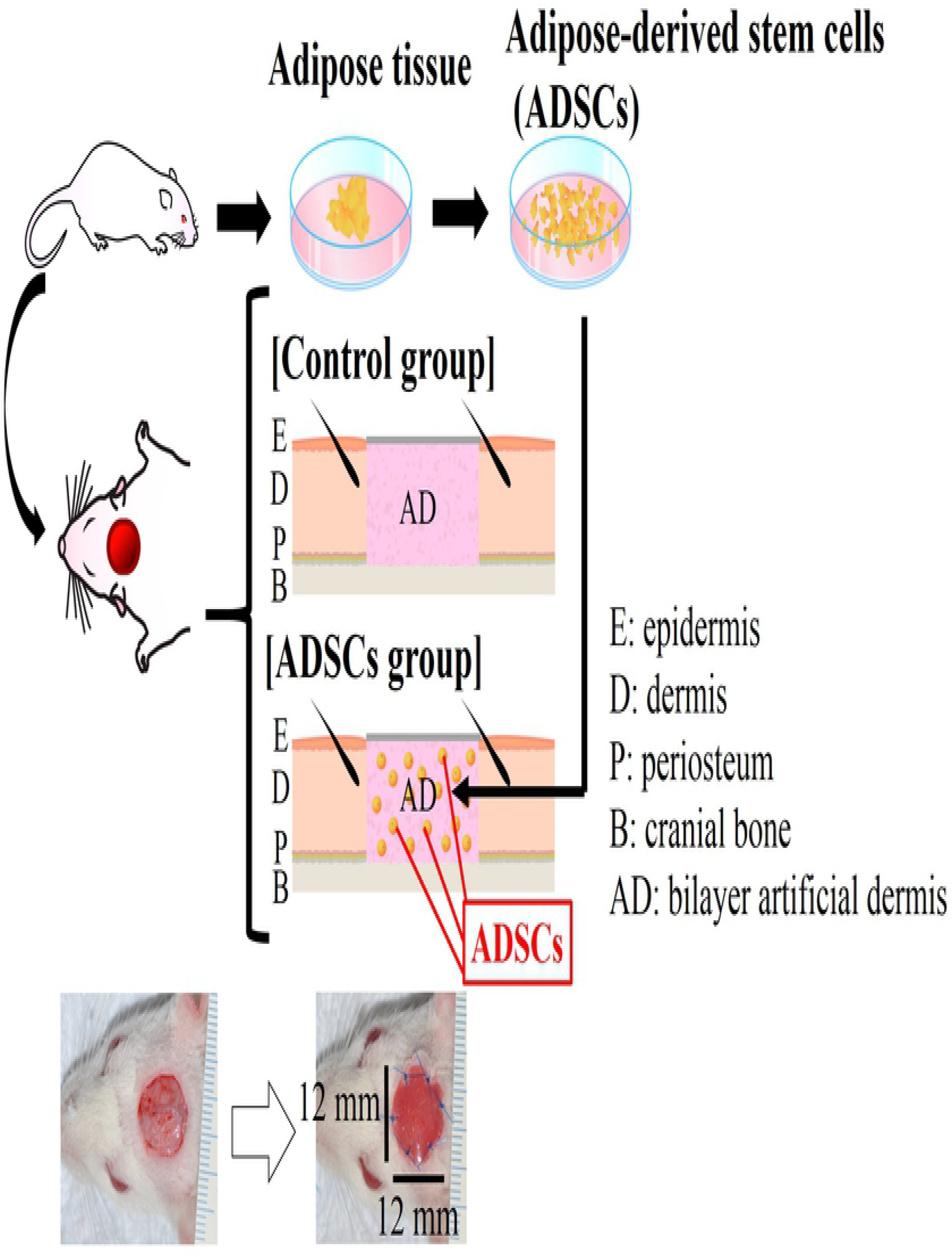
Schematic representation of the experimental procedure for transplanting autogenic ADSCs with an artificial dermis into a rat wound model with exposed bone. Scale bar: 1 mm.

### Artificial dermis

We used the commercially available artificial dermis Integra (Life Sciences Corp., Plainsboro, NJ, USA) for this experiment. Integra is widely used in the clinical treatment of deep partial-thickness and full-thickness burn wounds [7,8]. In addition, in clinical settings, Integra has been successfully used to promote healing of wounds with exposed bone [1,2].

### Isolation of ADSCs

ADSCs were isolated from the bilateral inguinal adipose tissue of the rats following a modification of a previously reported method [9]. In brief, the adipose tissue was washed with phosphate-buffered saline (PBS; Fujifilm Wako Pure Chemical Corporation, Osaka, Japan) and cut into strips. Collagenase (Fujifilm Wako Pure Chemical Corporation, Osaka, Japan) was dissolved in 20 mL of PBS to a final concentration of 0.1% to digest the adipose tissue for 60 minutes in a 37°C water bath (the mixture was shaken every 15 minutes during the digestion period). Immediately after the reaction was completed, 20 mL of Dulbecco’s modified Eagle’s medium (DMEM; Fujifilm Wako Pure Chemical Corporation, Osaka, Japan) was added to neutralize the collagenase activity. The resulting solution was then filtered by a 40-μm cell strainer (Corning Inc., Corning, NY, USA), the filtrate was centrifuged at 700×g for 6 minutes at 25°C, and the supernatant was removed. The remaining deposit was added to DMEM supplemented with 10% fetal bovine serum (FBS; Corning Inc., Corning, NY, USA) and 1% penicillin/streptomycin (P/S; Fujifilm Wako Pure Chemical Corporation, Osaka, Japan), plated on a 60.1-cm^2^ tissue culture dish (TPP Techno Plastic Products, Trasadingen, Switzerland), and cultured at 37°C in a 5% CO_2_ incubator. After 24 hours, the debris was removed by washing with PBS, and fresh medium was added. The ADSCs were selected based on the ability of cells to adhere to the culture plate. The cells were passaged with 0.25% trypsin-ethylenediaminetetraacetic acid (Fujifilm Wako Pure Chemical Corporation, Osaka, Japan) on day 3 and transferred to a new dish. Cells were used at the third passage in all experiments. Kato et al. [10] proved that these cells have MSC-like self-renewal, adipogenesis, and osteogenesis properties.

### ADSC seeding

Integra was placed with the silicone side positioned downward. In this scaffold, 1.0 × 10^6^ ADSCs/mL were seeded drop-wise with 40 μL of DMEM supplemented with 10% FBS and 1% P/S according to the fluid capacity of the scaffold. The scaffold of the control group was placed in a separate tissue culture dish with the same culture medium but without ADSCs. All dishes were incubated at 37°C with 5% CO_2_. After 24 hours, just prior to transplanting, the debris was removed by washing with PBS, and fresh medium was added.

### Wound area

On the day of surgery and at the established endpoints (3, 7, 14, and 21 days after surgery), the wound area was measured by tracing the wound margin on the photograph followed by calculation of the pixel data using ImageJ software (National Institutes of Health, Bethesda, MD, USA). The global wound area (%) was calculated according to the residual wound area on a given day (tx) relative to the wound area measured on the day of surgery, as follows:

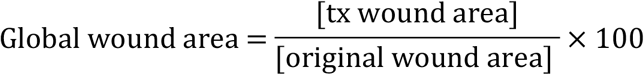

### Histological analysis of paraffin-embedded tissues

At the established endpoints (3, 7, 14, and 21 days after surgery), the wounds were harvested with approximately 5-mm margins and the cranial bone attached (n = 5 at each time point). A part (right quarter) of the wound was used for subsequent real-rime RT-PCR analysis. The remaining part of the wound was fixed in 10% neutralized formalin solution and dehydrated using an ethanol gradient (70%, 80%, 90%, and 100%). The fixed specimens were decalcified in 10% formic sodium citrate solution, embedded in paraffin, and sectioned in the coronal plane. The sections were stained with hematoxylin and eosin and Masson’s trichrome, and the slides were observed using an optical microscope (Biorevo BZ-9000; Keyence Co., Osaka, Japan).

### Immunohistochemistry

Tissue samples were embedded in paraffin and sectioned in the coronal plane, and the blood vessel density was analyzed immunohistochemically (n = 5 at 14 days after surgery per group). Blood vessel endothelial cells were immunohistochemically stained with an anti-CD31 antibody (Abcam, Cambridge, Britain, ab28364, 1:50). The sections were incubated in Liberate Antibody Binding Solution (LAB solution; Polysciences, Philadelphia, PA, USA) at room temperature for 15 minutes, and then Protein Block Serum-Free (Dako, Glostrup, Denmark) and phosphate buffer containing hydrogen peroxide (Peroxidase-Blocking Solution, Dako, Glostrup, Denmark) were added to the sections for 10 minutes each for blocking. Primary anti-CD31 antibody in Tris-HCl buffer containing stabilizing protein and 0.015 mol/L sodium azide (Dako Antibody Diluent, Dako, Glostrup, Denmark) were then added to the slides, which were incubated at 4°C overnight. After addition of the secondary antibody, Dako REAL^TM^ EnVision^TM^/HRP, Rabbit/Mouse (ENV, Dako, Glostrup, Denmark), the slides were further incubated for 30 minutes at room temperature. The samples were developed with DAB chromogen (Dako REAL^TM^ DAB+ Chromogen, Dako, Glostrup, Denmark) and the substrate and observed under the microscope. Once the desired signal-to-noise ratio was achieved, the reaction was stopped by washing the slides in deionized water. To quantify vascularization within the wound, the blood vessel densities of five animals in each group at 14 days after surgery were determined by measuring the vascular area in the wound area of the specimen.

The blood vessel densities were quantified at the established endpoints in the immunohistochemically stained samples. Quantification was performed in five randomly selected high-power fields of five non-consecutive tissue sections per wound within each group [11]. Only vessels with a diameter < 50 μm [12] were considered for this analysis. In this study, it was inevitable that the collagen of the artificial dermis would become stained with an anti-CD31 antibody. Therefore, we excluded this area for measurement of the vascular area. The vessel area in the selected field of each specimen was observed with a fluorescence microscope (U-RFL-T; Olympus, Tokyo, Japan), and the images were analyzed with ImageJ software (National Institutes of Health, Bethesda, MD, USA). The percentage of the relative area of CD31-positive vessels was calculated using ImageJ software from the following equation for each time point (tx):

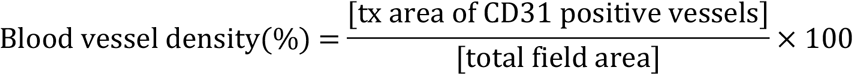

### DiI labeling

To confirm survival potential and location of the transplanted ADSCs, the ADSCs were labeled with the fluorescent dye DiI (Vybrant^®^ DiI Cell Labeling Solution; Life Technologies, Carlsbad, CA, USA) prior to transplantation. DiI binds to cellular thiols and has long-term stability, which enables the tracing of DiI-labeled transplanted cells in the host tissue. This experiment involved a separate group of rats (n = 3). The concentration of ADSCs was adjusted to 1.0 × 10^6^ cells/mL, and 5 μL/mL of DiI was dissolved in this medium and incubated for 15 minutes at 37°C in a 5% CO_2_ incubator for ADSC labeling. After the reaction was completed, the filtrate was centrifuged at 1000 rpm for 5 minutes at 25°C, and the supernatant was removed. Once the DiI was completely removed from the filtrate, ADSCs were centrifuged twice with DMEM under the same setting, and the supernatant was removed. DiI-labeled ADSCs were seeded to the Integra scaffold and transplanted as described above. At day 21 after transplantation of the labeled ADSCs, a frozen section was prepared using Kawamoto’s film method [13] in the coronal plane. The survival of the transplanted cells was then determined in the unstained samples.

### Real-time RT-PCR

RNA was extracted from the granulation tissue of the rats using a NucleoSpin^®^ RNA II kit (Takara Bio, Otsu, Japan). Each sample was harvested from the right quarter of a wound with approximately 5-mm margins without the cranial bone, and was disrupted and homogenized using a syringe (n = 5 at each time point per group). The absorbance of the resulting total RNA concentrations was determined on an ultraviolet-visible spectrophotometer with absorbance read at 260/280 nm (NanoDrop Lite; Thermo Scientific, Waltham, MA, USA). For real-time RT-PCR, 6 μg of mRNA was reverse-transcribed with RevertAid First-Strand cDNA Synthesis Kit (Thermo Scientific, USA) using a thermal cycler (T100TM Thermal Cycler; Bio-Rad, Hercules, CA, USA). Real-time RT-PCR was then carried out on an ABI Prism 7900 apparatus (Applied Biosystems, Foster City, CA, USA) using Sybr Green PCR Master Mix (Applied Biosystems, Foster City, CA, USA) per the manufacturer’s instructions. Primers for rat glyceraldehyde-3-phosphate dehydrogenase (*Gapdh*), basic fibroblast growth factor (*Fgfb*), vascular endothelial growth factor (*Vegf*), transforming growth factor beta 1 (*Tgfb1*), and beta 3 (*Tgfb3*) were purchased from Hokkaido System Science Co., Ltd. Japan; the sequences are listed in Table 1. The amplification parameters were an initial 95°C incubation step for 15 minutes, followed by 20 amplification cycles of 94°C for 15 seconds, 60°C for 30 seconds, and 72°C for 30 seconds. The reactions ended with a 72°C extension step for 7 minutes, followed by storage at 4°C overnight. The expression levels of each target gene were calculated relative to the level of *Gapdh* for each sample.

**Table 1.**
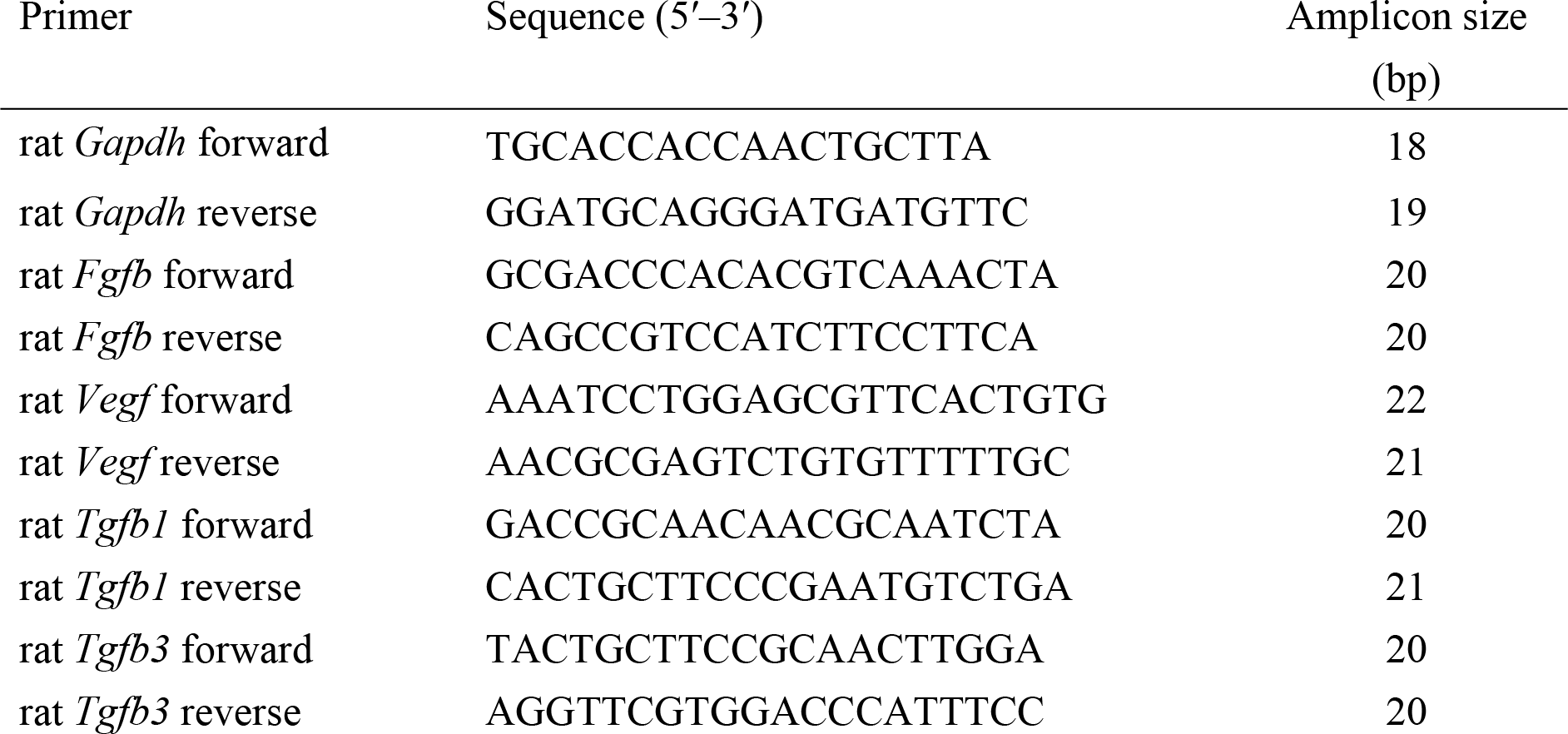
Sequences of primers used in real-time RT-PCR amplification.

## Statistical analysis

All statistical analyses were performed using the Statistical Package for Social Sciences version 23.0 (SPSS, Inc., Chicago, IL, USA). All results are presented as the mean ± standard deviation (SD). The data were normally distributed; comparisons between two groups (i.e., control vs. ADSCs) in the global wound area, the percentage of the relative area of CD31-positive vessels, and relative gene expression levels were performed using Student’s *t*-test; *p* < 0.05 was considered statistically significant.

## Results

### Global wound area

Digital photographs were obtained immediately following surgery and at 3, 7, 14, and 21 days after surgery (Fig. 2a). The average global wound area was significantly smaller in the ADSC group than in the control group on days 3, 7, and 14 after surgery (Fig.2b). However, there was no significant difference between the two groups at day 21 because the wounds in almost all the rats were completely cured at that point (Fig. 2b).

**Fig. 2.**
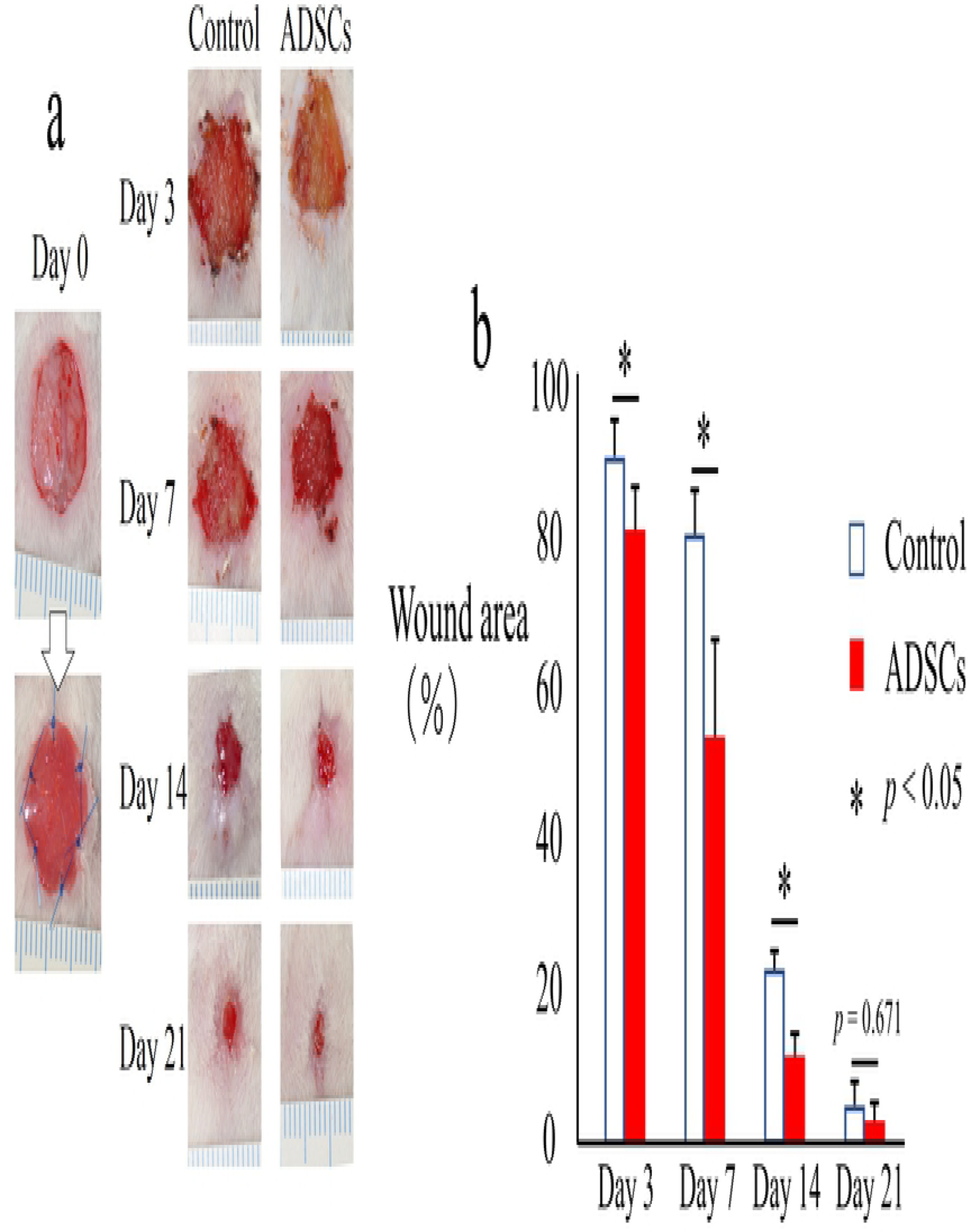
Images of wounds with exposed bone and comparison of healing with and without ADSC transplantation. (a) Macroscopic images of the wound with exposed bone. Scale bar: 1 mm. (b) Measurement and analysis of the wound area with or without ADSC transplantation. The data represent the mean ± SD.*p < 0.05.

## Histology

In magnified microphotographs of sagittal sections of hematoxylin and eosin-stained specimens collected 3 days after surgery (Fig. 3a, b), inflammatory cells, mainly neutrophils, as well as red blood cells were observed within the collagen sponges in both groups. The number of cells was higher in the ADSC group than in the control group. Seven days after the surgery (Fig. 3c, d), fibroblasts were observed in the collagen sponges with slight vascularization, and the fibroblast numbers and tissues density were higher in the ADSC group than in the control group. Fourteen days after the surgery (Fig. 3e, f), a thick and dense dermal layer was observed in both groups. At 21 days after the surgery (Fig. 3g, h), most of the wounds showed epithelization both macroscopically and histologically.

**Fig. 3.**
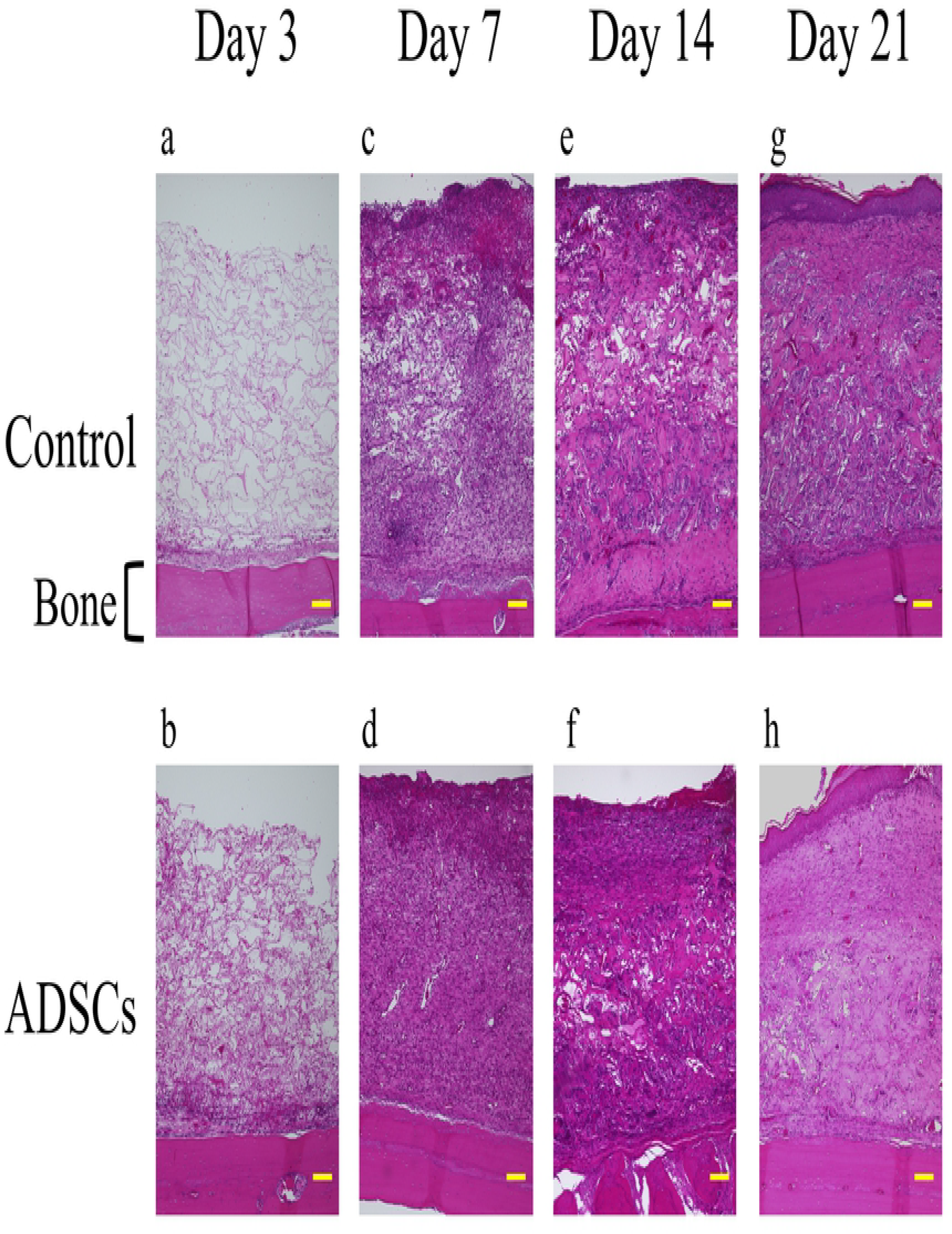
Representative magnified images (100x) of hematoxylin and eosin-stained histological sections of the center of the wound along with implantation time. Histological sections at day 3, 7, 14, and 21 highlight the wound healing progression along the experimental time frame. Scale bars indicate 100 μm for all panels.

In Masson’s trichrome-stained sections, especially at 7 and 14 days (Fig. 4c–f), higher-density granulation tissue was noted for the ADSC groups. In addition, the wounds of the ADSC groups showed greater tissue thickness and more homogeneous granulation tissue than those of the control groups at all time points (Fig. 4).

**Fig. 4.**
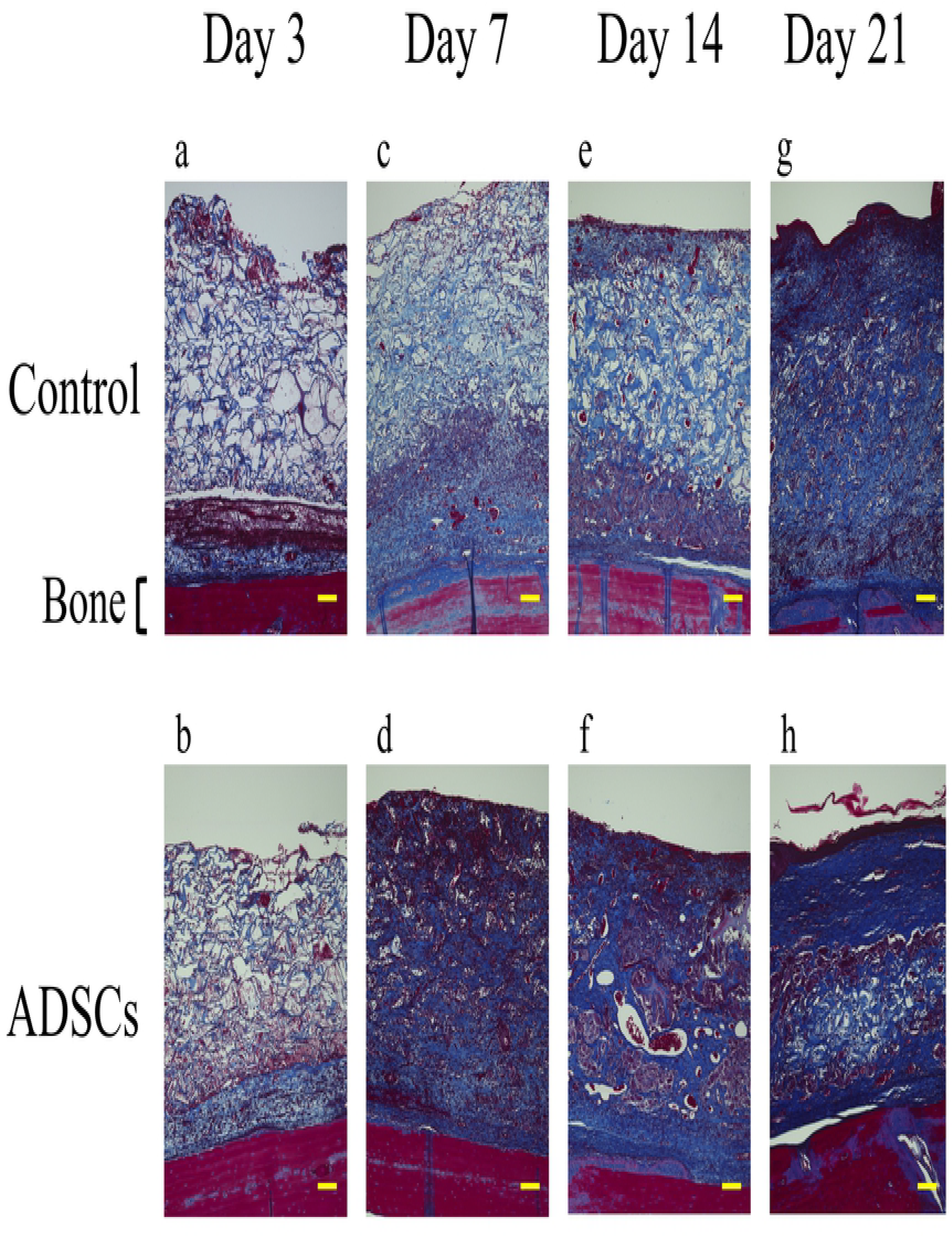
Representative magnified images (100x) of Masson’s trichrome-stained histological sections of the center of the wound along with implantation time. Histological sections at day 3, 7, 14, and 21 highlight the wound healing progression along the experimental time frame. Scale bars indicate 100 μm for all panels.

### Immunohistochemistry

The blood vessel density was quantified 14 days after surgery. The extent of neovascularization in the injured tissues was evaluated by immunostaining detection of CD31-expressing endothelial cells in paraffin-embedded tissue samples sectioned in the coronal plane (Fig. 5a~d). The blood vessel density in the wound was increased by 1.6-fold in the ADSC group compared with that in the control group after 14 days, representing a statistically significant difference (*p* < 0.01) (Fig. 5e).

**Fig. 5.**
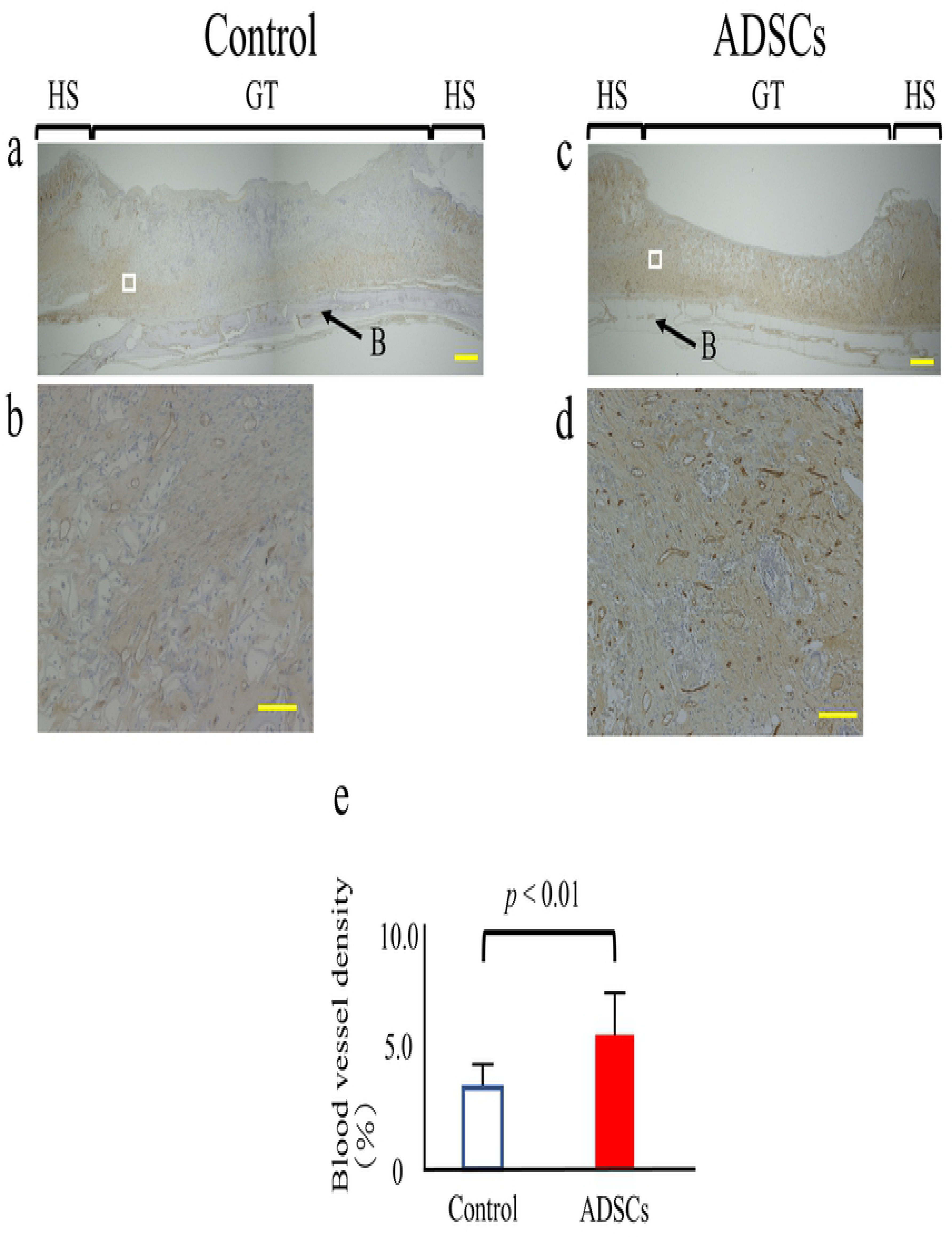
Quantitative analysis of wound neovascularization. (a~d) Control and ADSC specimens at 14 days after surgery were stained for CD31, a blood vessel endothelial cell marker (brown) (a, c: 40x magnification. Scale bars = 1 mm). For each group, the figure below is a micrograph of higher-magnification images of the boxed regions in the figure above (b, d: 400x magnification. Scale bars = 100 μm). (e) Blood vessel density was calculated by dividing the area of CD31-positive vessels by the total area. The data represent the mean ± SD. HS = healthy skin; GT = granulation tissue; B = cranial bone.

### DiI labeling

Magnified micrographs of sagittal sections of hematoxylin and eosin-stained specimens that were collected from the ADSC group 21 days after surgery are shown in Fig. 6a. Histologically, most wounds showed epithelization, and high-density granulation and homogeneous tissue were noted under the epithelization. For identification of tissues using DiI labeling, the gray scale values of the DiI-labeled section were used (Fig. 6b). In the DiI-labeled section, at 21 days after surgery, numerous DiI-positive (red) cells were distributed throughout the granulation tissue (Fig. 6c, d).

**Fig. 6.**
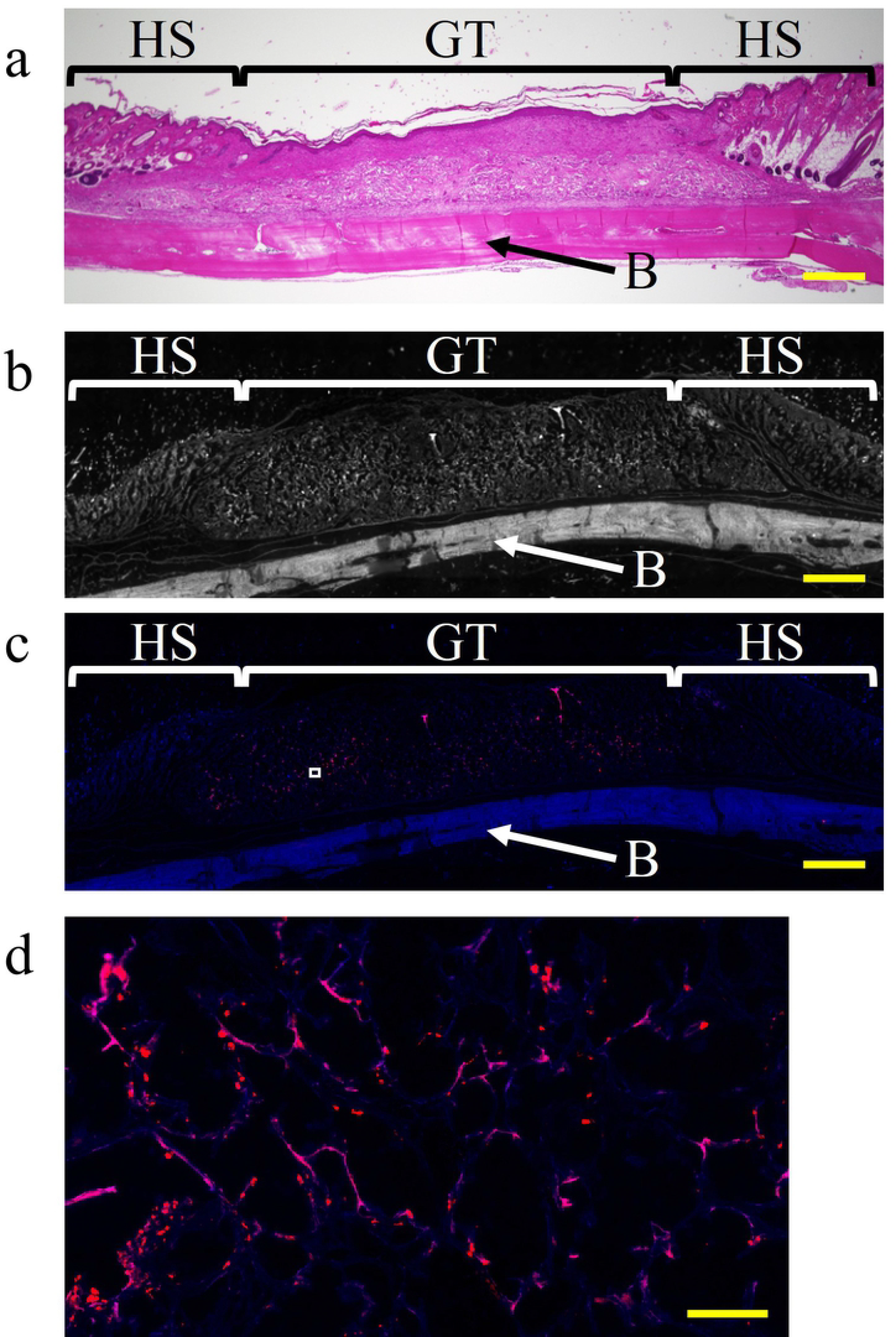
Representative images of DiI labeling of the center of the wound along with implantation time. Frozen sections were prepared at 21 days after transplantation of ADSCs labeled with DiI dye. (a) Magnified microphotographs of sagittal sections of hematoxylin and eosin-stained specimens collected from the ADSC group 21 days after surgery. (b) Grayscale values of frozen sections of each tissue (40× magnification); these values were similar to those for the frozen section in (c). Scale bar, 1 mm. (c) Frozen section with DiI labeling (40× magnification. Scale bar, 1 mm). (d) high-magnification image (500×) of the boxed region in (c). Scale bar, 100 μm. HS = healthy skin; GT = granulation tissue; B = cranial bone.

### Real-time RT-PCR

The ability of the transplanted ADSCs to promote neovascularization at the molecular level was assessed based on the expression of *Fgfb* and *Vegf* using real-time RT-PCR. Significantly higher expression levels of both genes were detected at all time points for the ADSC group than for the control group *(p* < 0.05) (Fig. 7).

**Fig. 7.**
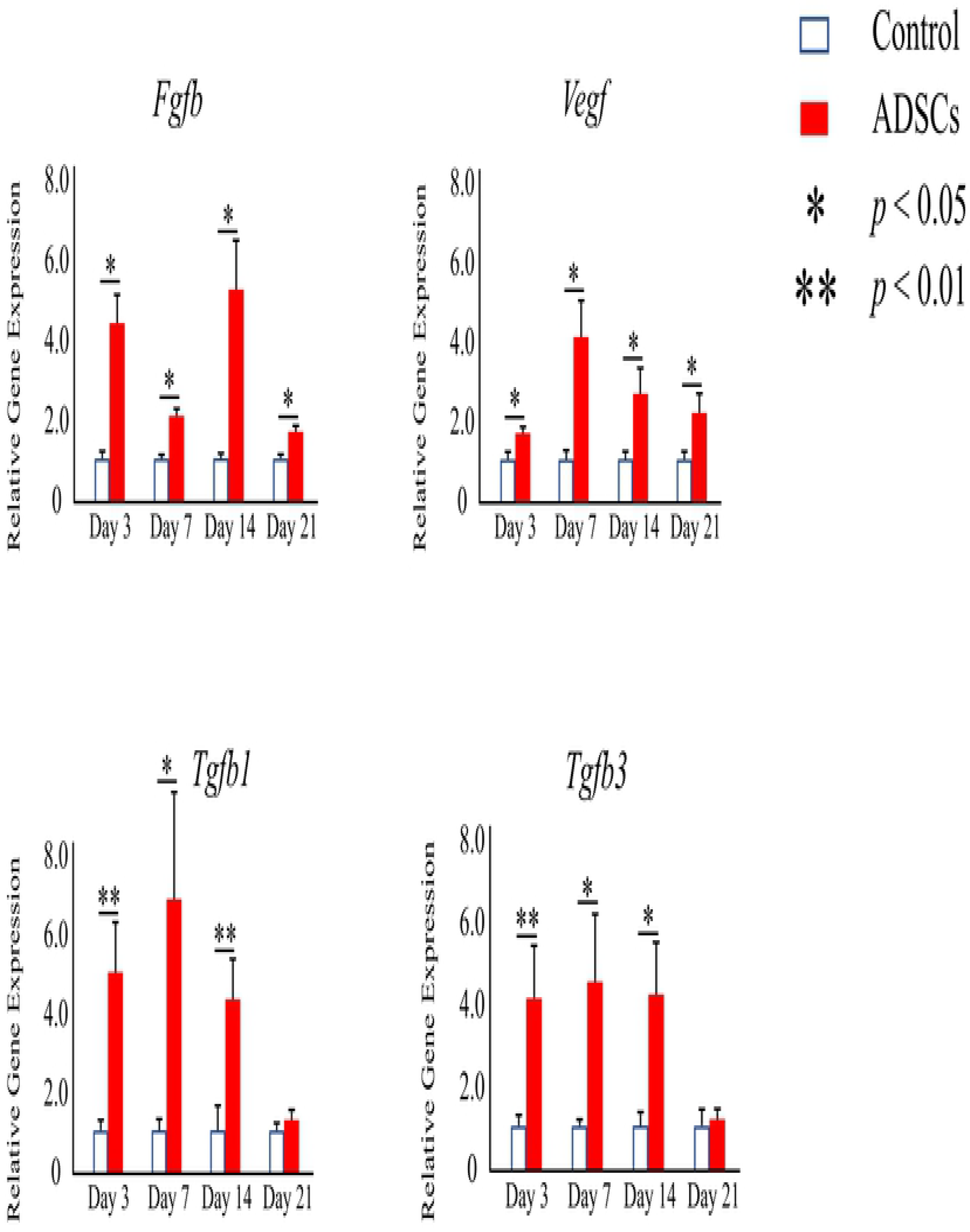
Relative expression levels of *Fgfb*, *Vegf*, *Tgfb1*, and *Tgfb3* among the different groups at different time points of implantation determined using real-time RT-PCR. The data represent the mean ± SD. *\*p* < 0.05, *\*\*p* < 0.01.

Moreover, the expression levels of *Tgfb1* and *Tgfb3* were examined to determine the potential biomechanical effect of ADSCs that could affect extracellular matrix deposition, organization, and scarring. Significantly higher expression levels of both genes were detected in the ADSC group until 14 days after surgery than in the control group (Fig. 7).

## Discussion

This study is the first report, to our knowledge, of using autogenic ADSCs with artificial dermis to heal wounds with exposed bone. The regeneration and introduction of capillaries inside the wound are important to wound healing because tissue regeneration commonly requires blood flow to supply oxygen and nutrition and remove waste. However, wounds with exposed bone are strongly deficient in terms of blood flow. In this study, we demonstrated that ADSCs increased the blood vessel density. Data from many studies have indicated that ADSCs exhibit improved and more rapid treatment effects in ischemic wounds such as burn wounds [14,15], radiation ulcers [16], or diabetic wounds [10,17]. Consistent with these previous findings, we showed that the wound area decreased significantly earlier when ADSCs were applied to wounds with exposed bone. The increased blood vessel density resulting from ADSCs at the ischemic wound site may help promote wound healing in cases with exposed bone.

The contributions of ADSCs to the complex wound-repair processes, which comprises inflammation, granulation, and remodeling, have been documented [18,19]. ADSCs have the potential to differentiate into several cell types (including endothelial cells), secrete angiogenic and anti-apoptotic factors [20,21], and exhibit various advantageous properties (such as paracrine activity and angiogenic potential) [22,23]. By DiI labeling, we showed the continued presence of ADSCs in granulation tissue up to 21 days post-transplantation in wounds with exposed bone. The results might not be sufficient to state that ADSCs directly differentiated into endothelial cells because we could not clearly check the form, nature, or detailed locations of the DiI-positive (red) cells.

However, numerous DiI-positive (red) cells were distributed throughout the granulation tissue (Fig. 4c). This result suggests that ADSCs might differentiate into other cell types, such as fibroblasts or blood vessel endothelial cells. Moreover, in terms of paracrine activity and angiogenic potential, the significantly higher expression levels of *Fgfb* and *Vegf* in the ADSC group than in the control group elicited improved neovascularization in the ADSC group. *Fgfb* and *Vegf* are known to play important roles during wound healing and can be secreted by ADSCs, thus influencing neovascularization [24,25]. In a previous study, significantly higher expression of *Fgfb* and *Vegf* were detected for a group with ADSCs added to a full-thickness excisional wound [26].

In full-thickness excisional wounds (without exposed bone), ADSCs exhibited increased *Tgfb3* expression, thereby decreasing the scar size and facilitating better collagen organization, scar pliability, and a more mature collagen arrangement [25,27]. Further, Zonari et al. [26] reported that the use of artificial dermis with ADSCs reduced *Tgfb1* expression in full-thickness excisional wounds. *Tgfb3* was previous found to promote normal collagen organization [25,28]. However, *Tgfb1* overexpression resulted in excessive fibroblast migration, myofibroblast differentiation, and scar formation [29]. In this study, significantly higher expression levels of both *Tgfb1* and *Tgfb3* were observed in the ADSC group than in the control group. We believe that, in wounds with exposed bone, higher expression levels of both Tgfb1 and Tgfb3 are preferable for covering the bone surface. Our data demonstrated that ADSCs exhibited increased expression of both *Tgfb1* and *Tgfb3* in wounds with exposed bone and promoted both scar formation and normal collagen organization for covering the bone surface. The main limitation of this study is that wound-healing mechanisms are different between humans and rodents. Humans display re-epithelialization and granulation tissue formation–based wound healing, whereas rats or mice exhibit contraction-based wound healing. Therefore, in a rodent model of full-thickness excisional wound of the back, some studies use silicone rings around the wound to reduce contraction upon wounding, thus allowing a system more representative of human wound healing [18,30]. However, in our model, using silicone rings is difficult; moreover, they result in serious stress to the model. In this study, artificial dermis was placed on a wound and fixed with nylon threads to reduce wound contraction and prevent wound enlargement as a result of the loose skin of the rats.

## Conclusions

In this study, to facilitate clinical application, we used ADSCs that were simply seeded drop-wise with a medium, along with the artificial dermis Integra that is commonly used as a scaffold. Further, in this study, we used autogenic ADSCs that have several important advantages regarding the convenience of their clinical application and fewer ethical concerns compared with those for allogeneic or xenogeneic ADSCs. Although a potential limitation to the translational use of ADSCs in patients is the enzymatic isolation technique used, we think that this technology is highly clinically translatable since both ADSCs and artificial dermis are relatively easy to obtain. Our study represents the first report that an existing artificial dermis can maintain autogenetic ADSCs, which can promote the vascularization capacity and enhance wound healing in a wound with exposed bone.

## Acknowledgments

We thank Ms. Yoko Kasai for her skillful technical assistance in the immunohistochemical analysis of paraffin-embedded tissues.

